# Single-cell characterization and machine learning-based classification reveal the transcriptional identity and antigen-experienced features of peripheral CD4⁺CD8⁺ double-positive T cells

**DOI:** 10.64898/2026.01.01.697324

**Authors:** Eunji Shin, Seung Gyu Yun, Yunjung Cho

**Affiliations:** Department of Laboratory Medicine, Korea University Anam Hospital, Seoul, Republic of Korea; Department of Laboratory Medicine, Korea University College of Medicine, Seoul, Republic of Korea

**Keywords:** Double Positive T cells, classification, machine learning, single-cell RNA sequencing, single-cell T cell receptor sequencing

## Abstract

T cell mediated immunity depends on diverse subsets with distinct regulatory and cytotoxic roles. While CD4⁺ and CD8⁺ T cells have traditionally been viewed as separate lineages, double-positive T (DPT) cells coexpressing both markers have emerged as a rare subset with potential roles in immune modulation, cytotoxicity, and memory. Here, we integrated single-cell RNA sequencing, single-cell TCR sequencing, and supervised machine learning to characterize transcriptionally defined DPT cells in human peripheral blood. DPT cells formed a reproducible transcriptional state distinct from conventional single-positive T cells and exhibited gene expression programs associated with cytotoxic effector potential, antigen experience, and immune cell migration. Clonal repertoire analysis revealed enrichment of expanded clonotypes and sharing of identical TCR clonotypes with both CD4⁺ and CD8⁺ T cell populations, indicating shared antigen-driven clonal histories rather than a distinct developmental lineage. To enable robust identification of this rare state, we developed a machine learning classifier trained on sorted CD4⁺ and CD8⁺ T cells, which outperformed marker-based approaches. Application of this framework to public COVID-19 single-cell datasets demonstrated its generalizability under immune perturbation. Together, these findings establish DPT cells as a reproducible, antigen-experienced transcriptional state with cytotoxicity- and migration-associated programs, and provide a framework for systematic identification of rare T cell states, expanding our understanding of T cell diversity.

## Introduction

T cells are central components of the adaptive immune system, orchestrating antigen-specific immune responses and establishing immunological memory [1]. Conventional CD4⁺ helper T cells and CD8⁺ cytotoxic T cells represent the two major single-positive T cell lineages that coordinate immune regulation and targeted elimination of infected or malignant cells [2]. These functions are mediated through the T cell receptor (TCR), whose vast diversity, generated by somatic recombination, enables precise recognition of a wide array of antigens. Upon antigen encounter, T cells undergo clonal expansion and differentiation, giving rise to effector populations and long-lived memory cells that collectively ensure durable immune protection [3].

Beyond these well-characterized single-positive T cell populations, accumulating evidence has highlighted the presence of unconventional T cell subsets, including CD4⁺CD8⁺ double-positive T (DPT) cells, in peripheral blood and tissues [4–7]. Although double-positive T cells are classically regarded as a transient developmental stage within the thymus, multiple studies have reported mature DPT cells in the periphery under both physiological and pathological conditions [7,8]. Peripheral DPT cells have been associated with diverse immunological contexts, including infections [6,9,10], tumor surveillance [8,11], autoimmunity [12–14], and immune aging, and have been proposed to exhibit cytotoxic [6,8,15], immunomodulatory [5,16], or memory-related properties [17,18]. However, their rarity, phenotypic ambiguity, and overlap with canonical CD4⁺ and CD8⁺ T cell markers have hindered systematic characterization and have raised persistent questions regarding their origin, stability, and functional identity.

From a developmental perspective, T cell lineage commitment in the thymus is governed by a tightly regulated transcriptional program centered on the mutually antagonistic transcription factors *ThPOK* and *Runx3* [15,19–22]. *ThPOK* enforces CD4⁺ lineage identity by suppressing CD8-associated programs, whereas *Runx3* promotes CD8⁺ differentiation by repressing CD4 expression. The detection of mature peripheral T cells that transcriptionally coexpress CD4 and CD8 challenges this canonical dichotomy and suggests the existence of non-canonical transcriptional states or functional plasticity beyond thymic development. Dissecting whether peripheral DPT cells represent transient activation states, reprogrammed memory populations, or distinct functional subsets remains an open question.

Recent advances in single-cell RNA sequencing (scRNA-seq) and single-cell TCR sequencing (scTCR-seq) have enabled high-resolution interrogation of immune cell populations, providing unprecedented opportunities to study rare and poorly defined populations such as DPT cells [23,24]. While scRNA-seq offers powerful resolution for profiling immune heterogeneity, its susceptibility to stochastic gene detection, background RNA signals, and multiplet capture can introduce ambiguity in the interpretation of marker coexpression, particularly in rare cell populations [25–27]. Moreover, conventional marker-based annotation strategies that rely on a small number of genes may fail to robustly capture rare transcriptional states, particularly when applied across datasets or experimental conditions [28–31].

In this study, we sought to address these challenges by integrating scRNA-seq and scTCR-seq analyses with supervised machine learning to identify and characterize transcriptionally defined DPT cells in human peripheral blood. We analyzed memory CD4⁺ and memory CD8⁺ T cells isolated by magnetic-activated cell sorting from a single healthy donor across four independent blood draws, enabling assessment of reproducibility across repeated sampling while minimizing inter-donor variability. By focusing on transcriptional programs rather than individual marker genes, we aimed to define DPT cells as a coherent and reproducible transcriptional state within the memory T cell compartment.

To overcome the limitations of manual annotation and marker-based classification, we developed a random forest-based classifier trained on CD4⁺ and CD8⁺ memory T cell populations. This approach leverages multi-gene expression patterns associated with T cell identity, activation, and differentiation, allowing more robust detection of rare subsets such as DPT cells in the presence of technical noise. In parallel, paired scTCR-seq data enabled us to examine clonal relationships and antigen experience, providing an orthogonal line of evidence to support transcriptional classification.

Finally, to explore the generalizability of this framework beyond healthy individuals, we applied our classifier to publicly available single-cell datasets from patients with Coronavirus Disease 2019 (COVID-19) [32,33]. SARS-CoV-2 infection is characterized by profound perturbations of the T cell compartment, including lymphopenia, altered trafficking, and dynamic clonal responses. These features provide a stringent test case for evaluating whether transcriptionally defined DPT cells can be consistently identified under disease-associated immune remodeling.

By combining single-cell transcriptomics, TCR repertoire analysis, and machine learning-based classification, this study establishes a framework for the robust identification of transcriptionally defined DPT cells and provides a systematic characterization of their molecular and clonal features. Our findings highlight DPT cells as a reproducible memory T cell state with distinct transcriptional programs and antigen-experienced properties, setting the stage for future investigations into their functional roles and immunological significance.

## Results

### We identified a distinct subset of CD4+CD8+ T cells in healthy individuals that exhibit cytotoxic and migratory transcriptional signatures

To delineate the heterogeneity of peripheral T cells in healthy individuals, we analyzed paired single-cell transcriptomic and TCR sequencing data from peripheral blood mononuclear cells. Samples were obtained from a single healthy donor across four independent blood draws. At each time point, memory CD4⁺ and memory CD8⁺ T cells were separately enriched using magnetic-activated cell sorting (MACS) prior to sequencing. After quality control and computational doublet filtering, the integrated dataset comprised memory T cells derived from all four sampling time points.

Unsupervised clustering using a shared nearest neighbor modularity optimization algorithm identified nine distinct T cell clusters, visualized by Uniform Manifold Approximation and Projection (UMAP) (Fig. 1a). These clusters clearly segregated canonical CD4⁺ and CD8⁺ T cell populations and showed strong concordance with transcriptional profiles derived from MACS-sorted CD4⁺ and CD8⁺ memory T cells, supporting the fidelity of the sorting strategy.

**Figure 1.**
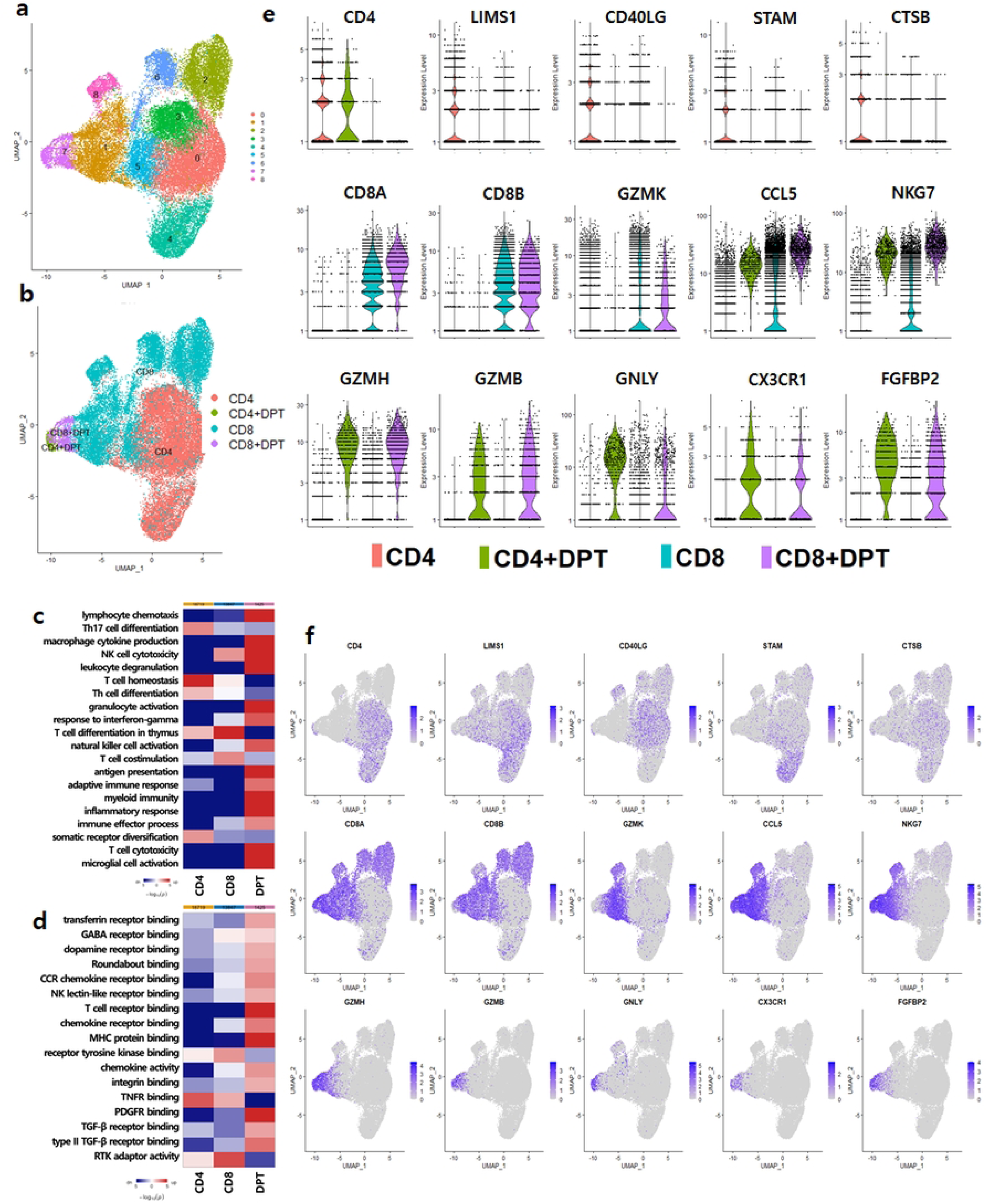
Molecular characterization of DPT Cells. **(a)** Overview of cell clusters in integrated single-cell transcriptomes from healthy donors, with cells colored according to nine clusters identified via SNN clustering, is presented. **(b)** UMAP projection of cells labeled CD4 (red), CD8 (blue), CD4+DPT (green), and CD8+DPT (purple). **(c-d)** The heatmaps show the results of gene set enrichment analysis, illustrating gene sets significantly enriched in (c) the immune-related GO category (GO:0002376) and (d) the binding-related GO category (GO:0005102) across distinct T cell populations. Red and blue indicate relative upregulation and downregulation, respectively, with color saturation corresponding to statistical significance (darker shades indicate greater significance). **(e)** Violin plots of marker genes from differentially expressed genes (DEGs) across subsets. (f) UMAP projection of T cells with coloring on the basis of the expression of marker genes.

Within this transcriptional landscape, we observed a subset of cells annotated as CD4⁺ T cells that localized within clusters predominantly composed of CD8⁺ T cells (Fig. 1b). These cells exhibited detectable expression of both *CD4* and *CD8A*/*CD8B* transcripts and displayed a gene expression profile distinct from that of conventional single-positive CD4⁺ T cells (S1 Table). We refer to these cells as double-positive T (DPT) cells, defined here as transcriptionally identified CD4⁺CD8⁺ candidates rather than a lineage-validated population.

Given the susceptibility of single-cell RNA sequencing to technical artifacts such as transcript drop-out, ambient RNA contamination, and doublets, multiple steps were taken to support the robustness of this identification. First, DPT cells did not exhibit elevated doublet scores compared with single-positive T cells. Second, DPT cells were reproducibly detected across all four independently collected blood samples, arguing against stochastic or batch-specific effects. Third, DPT cells formed a coherent transcriptional cluster characterized by shared gene expression programs, rather than occupying a continuous gradient between CD4⁺ and CD8⁺ single-positive populations. Together, these observations support the interpretation that DPT cells represent a reproducible transcriptional state within the memory T cell compartment.

Based on relative transcript abundance, DPT cells were further subdivided into CD4-derived DPT (CD4⁺DPT) and CD8-derived DPT (CD8⁺DPT) subsets, defined by predominant expression of *CD4* or *CD8* transcripts, respectively. Across the four sampling time points, CD4⁺DPT and CD8⁺DPT cells accounted for approximately 1 percent and 3 percent of the total memory T cell population. Although numerically rare, their consistent detection across repeated sampling suggests that these cells represent a stable transcriptional subset in peripheral blood.

Transcriptomic profiling revealed that DPT cells are characterized by a distinct gene expression program associated with cytotoxic effector potential and immune cell migration (Fig. 1c-f; S2 Table and S3 Table). At the transcript level, DPT cells showed elevated expression of multiple genes encoding cytotoxic effector molecules, including *GZMB*, *GZMH*, *GZMM*, and *GNLY*, as well as genes involved in granule exocytosis and proteolytic activity such as *NKG7*, *CST7*, and *CTSW*. These expression patterns are commonly associated with cytotoxic lymphocyte programs, but are interpreted here as indicative of cytotoxicity-associated transcriptional programs rather than direct evidence of functional cytolytic activity.

Consistent with this interpretation, gene set enrichment analysis demonstrated significant enrichment of immune effector related pathways, including T cell mediated cytotoxicity, natural killer cell mediated cytotoxicity, and leukocyte degranulation (Fig. 1c; S3 Table). In parallel, DPT cells expressed a distinct set of chemokines and receptors involved in immune cell trafficking and tissue surveillance, including *CCL4*, *CCL5*, *XCL1*, *XCL2*, and *CX3CR1*. Pathway enrichment analyses further highlighted biological processes such as chemokine receptor binding, lymphocyte chemotaxis, and granulocyte activation (Fig. 1c-d; S3 Table).

Collectively, these data indicate that transcriptionally defined DPT cells represent a reproducible memory T cell subset characterized by cytotoxicity-associated and migratory gene expression programs. While these findings suggest that DPT cells may be poised for effector and tissue-infiltrative functions, direct functional validation at the protein and cellular levels will be required to establish their cytolytic activity and immunological roles.

### Machine learning enables robust identification of rare DPT transcriptional states

Accurate identification of rare transcriptional states such as DPT cells is challenging when relying solely on individual marker genes, particularly in the presence of transcript drop-out. To address this limitation, we developed a supervised machine learning classifier based on a random forest algorithm trained on scRNA-seq data from MACS-sorted memory CD4⁺ and CD8⁺ T cells (Fig. 2a). The classifier was designed to integrate multi-gene expression patterns rather than rely exclusively on CD4 or CD8 transcript detection.

**Figure 2.**
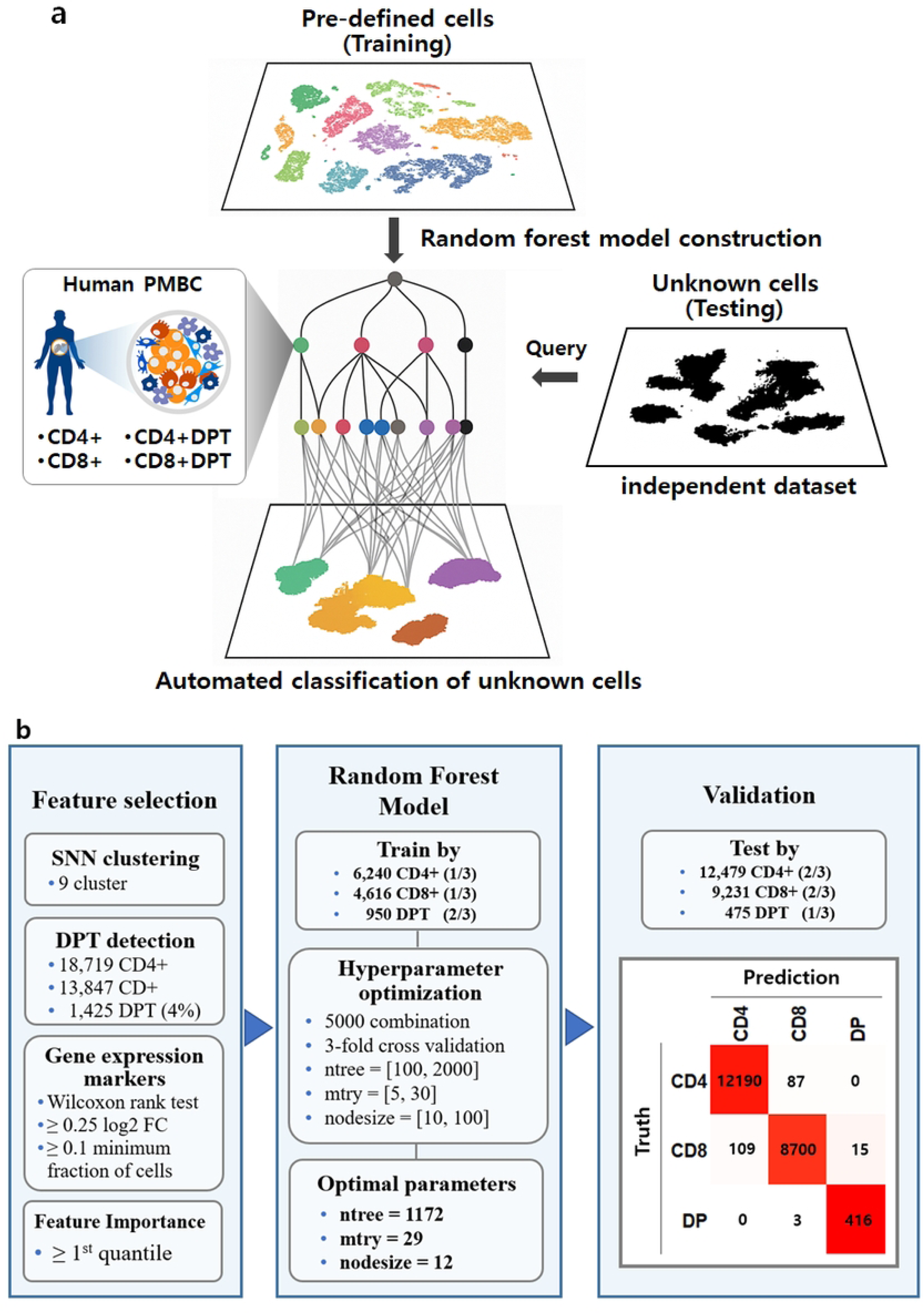
Development and validation of a supervised learning model for DPT cell classification. **(a)** Workflow of the machine learning model for T cell classification and prediction on independent datasets. **(b)** A schema showing the series of steps for creating a classification model, including feature selection, data splitting, hyperparameter tuning, and model validation.

Following hyperparameter optimization via grid search and undersampling to mitigate class imbalance, the final model consisted of 1,172 trees, with 29 variables considered at each split and a minimum terminal node size of 12. The classifier demonstrated strong generalization performance, with an out-of-bag (OOB) estimate of the overall error rate of 2.17%, indicating robust predictive stability without overfitting. Class-specific OOB error rates were 1.36% for CD4⁺, 2.73% for CD8⁺, and 4.74% for DPT cells, with the slightly elevated error for DPT cells likely reflecting their low prevalence and partial transcriptional overlap with SPT cell states.

Evaluation on the held-out test set confirmed the high discriminative power of the model, achieving an overall accuracy of 97.9% (95% CI: 97.7–98.1%) and a Cohen’s κ of 0.96, indicating near-perfect agreement beyond chance (Fig. 2b). The classifier maintained high sensitivity and specificity across all classes, including for the rare DPT population (sensitivity 96.8%, specificity 99.6%, balanced accuracy 98.2%). Notably, although the positive predictive value for DPT cells was lower than that of CD4⁺ or CD8⁺ cells, this behavior is expected in the context of severe class imbalance (∼2% prevalence) and underscores the intrinsic difficulty of precision in rare-cell classification tasks.

Importantly, the machine learning–based approach substantially outperformed a simple marker-based rule requiring concurrent detection of CD4 and CD8 transcripts, which was more susceptible to transcript drop-out and showed reduced recall for rare DPT cells. These results demonstrate that integrating multigene expression patterns provides a more reliable strategy for identifying DPT cells than reliance on individual marker genes alone.

Feature importance analysis revealed that, in addition to canonical markers (*CD4*, *CD8A*, *CD8B*), the classifier relied heavily on genes associated with cytotoxic programs (*CTSW, GNLY, GZMH, GZMK*) [34,35] and transcriptional regulators (*GATA3, FOXP3*) [36–38], as well as genes involved in metabolic and structural processes (*FTH1, PFN1, MALAT1, GABARAPL1*) [39–41] (S4 Table). Collectively, these findings indicate that the classifier captures a composite transcriptional signature reflective of the DPT state, providing a robust and generalizable framework for identifying this rare T-cell population beyond single-marker strategies.

### DPT cells show enhanced clonal sharing and expansion based on TCR repertoire analysis

To explore the antigen-specific nature of T cell-mediated adaptive immunity, we analyzed the distribution and dynamics of TCR clonotypes using scTCR-seq data from PBMCs. Clonotypes were defined based on 100% identical CDR3 amino acid sequences of paired TCR alpha (α) and beta (β) chains, and cells sharing the same α-β CDR3 sequence were considered identical clonotypes (Fig. 3a). Cells with detectable TRGV expression, corresponding to γδ T cells, were excluded from clonotype analyses to focus on αβ T cell-mediated responses. Using this definition, identical TCR clonotypes were quantified across DPT and SPT cell subsets.

**Figure 3.**
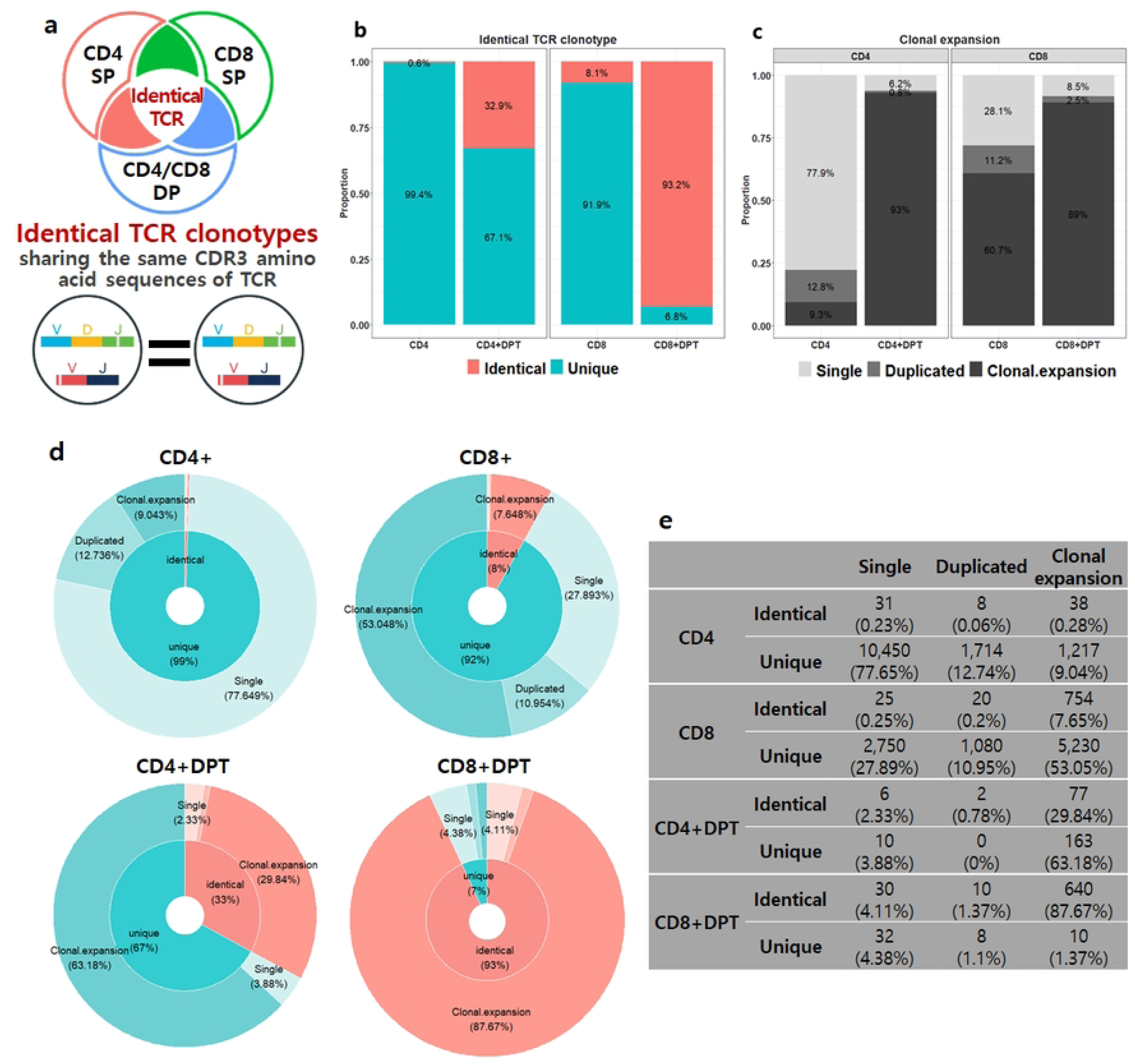
Clonal Expansion and TCR Diversity in T Cell subsets. **(a)** Illustration summarizing identical TCR clonotype. **(b)** Proportion of identical TCR clonotype for each subset. **(c)** Proportion of expanded CDR3 clones for each subset. **(d)** Donut charts displaying the combined results of identical TCR information and clonal expansion for all CDR3 clones. **(e)** A table presenting the combined results of identical TCR information and clonal expansion for all CDR3 clones, including counts and proportions.

Notably, multiple clonotypes were shared between DPT cells and either CD4⁺ or CD8⁺ SPT cells. Importantly, DPT cells exhibited a substantially higher frequency of these shared clonotypes compared with their single-positive counterparts (Fig. 3b), indicating that DPT cells likely arise from or coexist with conventional αβ T cells responding to common antigenic stimuli. We next assessed clonal expansion by examining the frequency distribution of individual clonotypes, defined as the number of cells sharing the same CDR3 sequence. Among conventional T cell subsets, CD8⁺ T cells showed greater clonal expansion than CD4⁺ T cells, consistent with their cytotoxic effector role. Remarkably, CD4⁺DPT cells displayed an approximately tenfold increase in clonal expansion relative to CD4⁺ SPT cells (Fig. 3c). When evaluating the entire repertoire of CDR3-defined clonotypes, DPT cells demonstrated both a higher number of shared clonotypes and a greater degree of clonal expansion compared with SPT cell subsets (Fig. 3d, e). Collectively, these results suggest that DPT cells represent a population of antigen-experienced αβ T cell clones that have undergone substantial proliferation.

### DPT cell subsets exhibit transcriptional features consistent with functional reprogramming and memory-like states

We next examined transcription factor expression and differentiation-associated features to contextualize the clonal enrichment observed in DPT cells (Fig. 4a-d). While *ThPOK* expression remained low across all subsets, *Runx3* expression was significantly elevated in both CD4⁺DPT and CD8⁺DPT cells relative to their single-positive counterparts (Fig. 4c, d). Notably, CD4⁺DPT cells expressed *Runx3* at levels exceeding those of conventional CD8⁺ T cells (Fig. 4d), suggesting that these cells are undergoing an active transition toward a CD8-like cytotoxic phenotype. This elevated *Runx3* expression implies functional reprogramming rather than lineage ambiguity, consistent with acquisition of effector functions.

**Figure 4.**
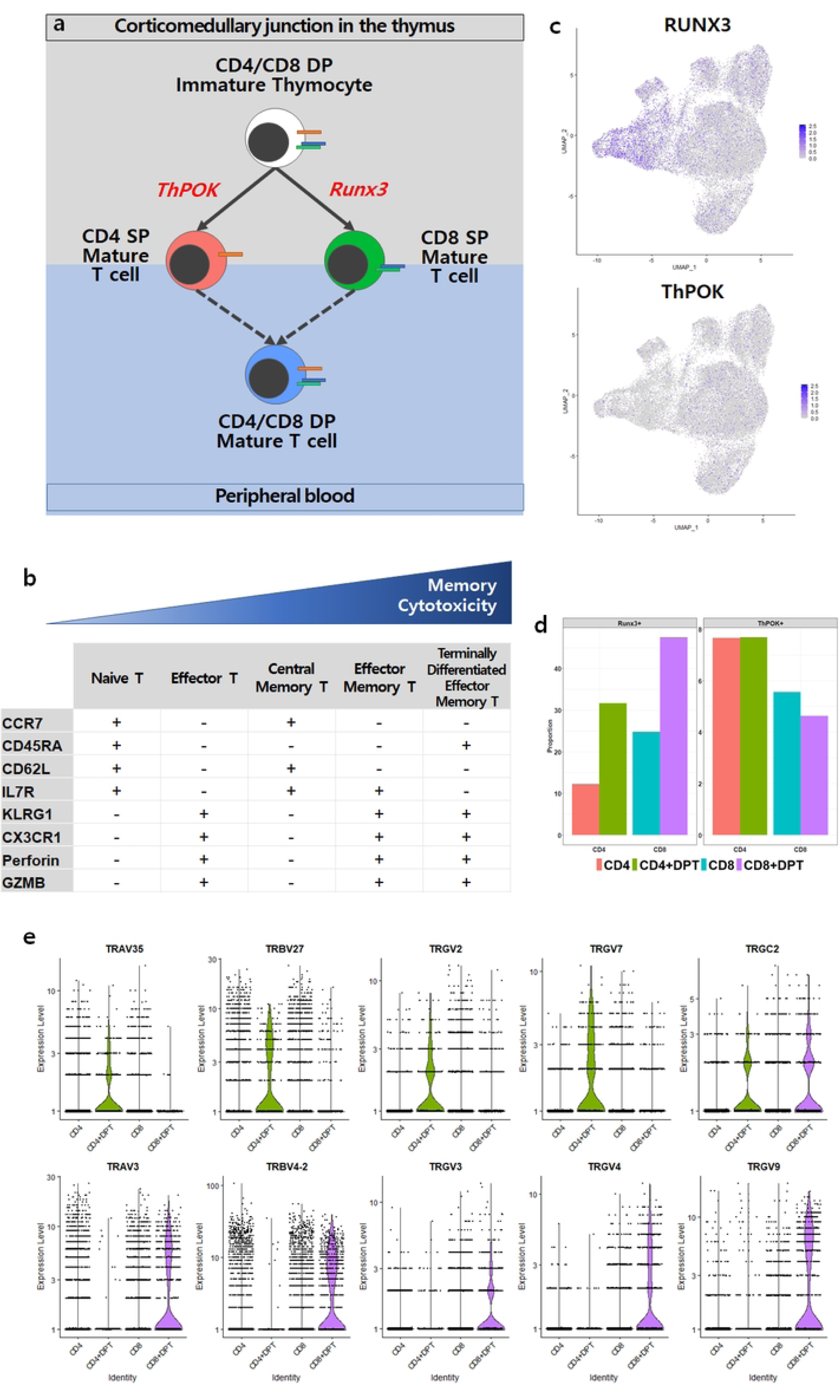
Regulation of human T cell development and differentiation by *ThPOK* and *Runx3* transcription factors. **(a)** Summary illustration depicting the potential regulation of T cell development and function by the *ThPOK* and *Runx3* transcription factors. **(b)** A conceptual framework describing the differentiation of human T cells. Naïve T cells undergo progressive differentiation into various populations of effector and memory cells following stimulation by specific antigens. **(c)** UMAP projection of expression levels for *Runx3* and *ThPOK*. **(d)** Proportion of genes upregulated by *Runx3* and *ThPOK*, stratified by median expression levels. **(e)** Violin plots illustrating the expression levels of genes associated with T-cell antigen recognition and signal transduction. **(f)** Density plots of gene expression associated with T cell differentiation. The median value is indicated by a line.

Analysis of TCR gene usage revealed distinct patterns between DPT subsets, with CD4⁺DPT cells enriched for *TRAV35* and *TRBV27*, and CD8⁺DPT cells preferentially expressing *TRAV3* and *TRBV4-2* (Fig. 4e). These differences suggest potential divergence in antigen recognition history between DPT subsets.

In addition, DPT cells displayed transcriptional features resembling terminally differentiated effector memory CD45RA⁺ T cells (TEMRA) (Fig. 4b, f), consistent with prior reports that mature DPT cells may persist long-term despite the short lifespan of immature DPT states [7,42]. Collectively, these findings support a model in which DPT cells represent an antigen-experienced, transcriptionally reprogrammed memory-like T cell state.

### The differential dynamics of DPT cells in COVID-19 provide insights into long-term immunity and disease resolution

To explore the relevance of DPT cells in viral infection, we applied our classifier to a publicly available single-cell dataset from COVID-19 patients (GSE158055) [33], selected for its longitudinal design, paired scRNA-seq and scTCR-seq data, and stratification by disease severity. The classifier maintained high accuracy (95%) in distinguishing CD4⁺ and CD8⁺ T cell subsets in this independent dataset (Fig. 5a), demonstrating cross-dataset generalizability.

**Figure 5.** Characterization of DPT Cells in COVID-19 Patients. **(a)** Confusion matrix employed to assess the performance of a classification model on an independent COVID-19 dataset. **(b)** Clonal expansion dynamics are depicted by the frequency of clonal expansion over time using the Generalized Additive Model (GAM) function. **(c)** Box plots illustrating the proportion of clonal expansion across each subtype.

Within the captured T cell population, CD4⁺DPT and CD8⁺DPT cells accounted for approximately 1% and 5% of cells, respectively. Given the known effects of lymphopenia and tissue trafficking during acute COVID-19, we interpret these values as relative proportions within sampled T cells, rather than as direct evidence of absolute expansion. Notably, predicted DPT cells in COVID-19 samples exhibited transcriptional and clonal features consistent with those identified in healthy individuals (S1 Fig).

Longitudinal analysis revealed distinct clonal dynamics associated with disease severity. In mild cases, DPT cells showed early clonal expansion followed by gradual contraction, whereas in severe cases, clonal expansion was attenuated during acute disease and partially recovered during convalescence (Fig. 5b, c). These patterns are consistent with a potential role for DPT cells in antiviral responses and immune reconstitution, although alternative explanations such as differential trafficking or survival cannot be excluded.

## Discussion

In this study, we provide a systematic characterization of transcriptionally defined CD4⁺CD8⁺ DPT cells within the human peripheral memory T cell compartment. By integrating single-cell transcriptomic and TCR repertoire analyses with supervised machine learning, we identify DPT cells as a reproducible and antigen-experienced T cell state that is consistently observed across repeated sampling from the same individual. Importantly, our findings frame DPT cells as a transcriptional state rather than a lineage-validated population, reflecting both the biological plasticity of mature T cells and the technical constraints inherent to single-cell profiling.

A central challenge in the study of peripheral DPT cells has been the ambiguity surrounding their identification. Apparent coexpression of *CD4* and *CD8* transcripts is especially vulnerable to technical artifacts such as transcript drop-out, ambient RNA contamination, and doublets [25–27], raising long-standing concerns about whether DPT cells reflect true biological states or technical noise. By integrating repeated sampling, stringent quality control, and multi-gene transcriptional analyses, our study provides evidence that DPT cells constitute a coherent and reproducible transcriptional state, rather than a stochastic artifact of single-cell sequencing.

Our transcriptomic analyses reveal that DPT cells are characterized by gene expression programs associated with cytotoxic effector potential and immune cell migration. Elevated expression of granzymes, granulysin, and genes involved in granule exocytosis, together with enrichment of pathways related to leukocyte degranulation and cytotoxicity, indicates that DPT cells are transcriptionally poised for effector-like activity [34,35,43]. Concurrently, the expression of chemokines and chemokine receptors linked to tissue trafficking suggests that these cells may be capable of localizing to inflamed or infected sites [44,45]. As these findings are based on transcriptional signatures and pathway enrichment analyses, future studies incorporating protein and cellular level analyses will be required to determine whether these transcriptional programs translate into active effector functions in vivo.

Clonal analyses further place DPT cells within the context of antigen-experienced T cell responses. The presence of expanded clonotypes and identical TCR clonotypes across DPT and SPT populations suggests that these cells arise from common antigen-driven clonal histories. These observations can be interpreted as suggesting that, following antigen-driven activation and clonal expansion, subsets of T cells diversify into distinct transcriptional and phenotypic states, one of which is characterized by the concurrent expression of *CD4* and *CD8*. The molecular signals and contextual cues that govern the emergence and maintenance of this state remain to be elucidated.

At the transcriptional regulatory level, DPT cells displayed low expression of *ThPOK* and elevated *Runx3* expression, a pattern that deviates from canonical T cell lineage commitment paradigms [7,20,21]. This configuration is consistent with transcriptional reprogramming toward a cytotoxic-associated state rather than incomplete lineage resolution. In addition, distinct patterns of TCR variable gene usage between CD4⁺DPT and CD8⁺DPT subsets suggest heterogeneity within the DPT compartment and raise the possibility that these subsets respond to different antigenic contexts. Together, these findings support the view that DPT cells are not simply intermediates between CD4⁺ and CD8⁺ lineages, but rather represent a distinct transcriptional adaptation within the memory T cell pool.

A key methodological contribution of this study is the application of supervised machine learning to the identification of rare T cell states. Conventional marker-based annotation strategies are particularly susceptible to transcript drop-out and often lack robustness across datasets or disease contexts. By leveraging multi-gene expression patterns, our classifier provides a more stable framework for identifying transcriptionally defined DPT cells, facilitating cross-dataset comparisons and reducing reliance on subjective manual annotation. In this sense, the classifier serves not only as a technical tool, but also as a conceptual framework for defining rare immune cell states beyond single-marker criteria.

Application of this framework to single-cell datasets from patients with Coronavirus Disease 2019 illustrates its generalizability under conditions of profound immune perturbation. DPT cells were identifiable in the context of SARS-CoV-2 infection and exhibited transcriptional and clonal features consistent with those observed in healthy conditions. However, given the profound lymphopenia, altered trafficking, and immune remodeling associated with COVID-19, changes in the relative abundance of DPT cells should be interpreted with caution. The observed severity-associated differences in clonal dynamics may reflect a combination of antigen-driven responses, differential survival, and tissue redistribution rather than direct expansion or contraction of the DPT population. Rather than establishing a direct clinical role for DPT cells in COVID-19, these analyses demonstrate the utility of our approach in tracking transcriptional states across physiological and pathological contexts.

Several limitations of our study should be acknowledged. First, our analyses are based on samples from a single donor, albeit collected across multiple independent time points. While this design enhances internal consistency and reduces inter-donor variability, it limits generalizability across individuals. Future studies incorporating larger and more diverse cohorts will be required to assess inter-individual variability. Second, our characterization of DPT cells is based primarily on transcriptomic and clonal features, and direct functional validation remains lacking. Protein-level assays, functional stimulation experiments, and cytotoxicity measurements will be necessary to determine whether the observed transcriptional programs correspond to effector function. Third, although clonal sharing implies antigen-driven activation, the specific antigens recognized by DPT cells remain unknown. Addressing these limitations will be essential for clarifying the biological roles and clinical relevance of this transcriptionally defined T cell state.

In summary, our study establishes transcriptionally defined DPT cells as a reproducible, antigen-experienced state within the human memory T cell compartment. By combining single-cell transcriptomics, TCR repertoire analysis, and machine learning-based classification, we provide a framework for the robust identification of rare T cell states and lay the groundwork for future studies aimed at elucidating the functional roles, developmental origins, and clinical relevance of DPT cells in health and disease.

## Methods

### Study Design

The aim of this study was to identify and characterize transcriptionally defined CD4⁺CD8⁺ double-positive T (DPT) cells in healthy human peripheral blood. Specifically, we sought to (i) define the transcriptional and TCR clonotypic features of DPT cells, (ii) examine their relationship to conventional single-positive T cell states, and (iii) develop a machine learning-based framework for robust classification of rare T cell populations.

Participants were prospectively recruited between 1 January 2017 and 1 March 2018. Peripheral blood samples were obtained from a single healthy donor across four independent blood draws, enabling assessment of reproducibility across repeated sampling while minimizing inter-donor variability. scRNA-seq and scTCR-seq were performed, followed by integrated bioinformatic analyses including quality control, transcriptional profiling, clonal tracking, and machine learning-based classification.

All procedures involving human participants were approved by the Institutional Review Board of Korea University Anam Hospital (IRB No. 2017AN0006), and written informed consent was obtained prior to study participation.

### Sample Collection and Preparation

PBMCs were isolated by Ficoll-Paque density gradient centrifugation. Memory CD4⁺ and memory CD8⁺ T cells were enriched separately using magnetic-activated cell sorting (MACS; Miltenyi Biotec) to reduce contamination from non-T cell populations. Sorting was performed according to the manufacturer’s protocol, yielding populations highly enriched for CD3⁺CD4⁺CD45RO⁺ and CD3⁺CD8⁺CD45RO⁺ T cells, respectively. Post-sorting transcriptional profiles confirmed the expected enrichment of lineage-specific markers.

Approximately 10,000 cells per sample were loaded for single-cell profiling using the Chromium Next GEM Single Cell Immune Profiling v2 Kit with Feature Barcode technology (10x Genomics). Gel beads-in-emulsion (GEM) generation, reverse transcription, cDNA amplification, and library preparation were carried out according to the manufacturer’s protocol provided in the Chromium Next GEM Single Cell Immune Profiling v2 User Guide (CG000331). The amplified cDNA was split to generate separate libraries for 5’ gene expression and TCR V(D)J profiling using the kit reagents.

### Single-Cell RNA Sequencing and TCR Sequencing

Final library quality and fragment size distribution were assessed using the Agilent TapeStation system. Sequencing was conducted on an Illumina NovaSeq 6000 platform with the following read configuration: Read 1, 26 bp; i7 index, 10 bp; i5 index, 10 bp; and Read 2, 90 bp. A sequencing depth of approximately 20,000 reads per cell was targeted for gene expression libraries and approximately 5,000 reads per cell for TCR libraries.

Barcoded cDNA libraries were demultiplexed into FASTQ files using the *mkfastq* pipeline from 10x Genomics Cell Ranger software (version 8.0). For gene expression analysis, reads were aligned to the GRCh38 reference genome, and barcode filtering and UMI counting were performed using the Cell Ranger *count* pipeline. TCR sequences were processed using the Cell Ranger *vdj* pipeline, and only productive, full-length sequences containing CDR3 regions were retained. Cells with matched TCR alpha (TRA) and beta (TRB) chains were selected for downstream TCR analysis. Cells expressing γδ TCR signatures were excluded by filtering out cells with detectable TRDC expression. Low-level detection of certain TRGV transcripts was observed in a subset of cells; however, these cells retained canonical αβ TCR expression and were therefore not classified as γδ T cells.

Quality control was conducted using the R package Seurat (version 5.1.0) [46]. Cells with fewer than 200 detected genes and with >5% mitochondrial gene expression were excluded. The data were normalized using the *NormalizeData* function, and variable features were identified using *FindVariableFeatures*. The data were then scaled using *ScaleData*, followed by principal component analysis (PCA). Dimensionality reduction was performed using UMAP via the *RunUMAP* function. Cell clustering was carried out using the *FindNeighbors* and *FindClusters* functions with a resolution parameter set to 0.4. Cluster-specific marker genes were identified using the *FindAllMarkers* function, and cell types were annotated by comparing cluster markers to canonical signature genes of known immune cell types.

### Identification of DPT Cells

DPT cells were operationally defined as transcriptionally identified T cells exhibiting detectable expression of both CD4 and CD8A/CD8B transcripts after quality control and doublet filtering. These cells formed a coherent transcriptional cluster distinct from conventional CD4⁺ and CD8⁺ SPT cells. Based on relative expression levels, DPT cells were further classified into CD4-derived (CD4⁺DPT) and CD8-derived (CD8⁺DPT) subsets.

### Functional Analysis

To characterize the molecular features of these DPT subsets, we performed differential gene expression analysis between DPT cells and SPT cell populations using the *FindMarkers* function in Seurat, which implements the Wilcoxon rank-sum test. P-values were adjusted for multiple comparisons using the Benjamini–Hochberg correction, and genes with an adjusted p-value < 0.05 were considered significantly differentially expressed.

To gain insights into the biological roles of DPT cells, we performed gene set enrichment analysis (GSEA) using the *CMScaller* R package [47]. Genes were ranked based on the product of log₂ fold change and the negative logarithm of the false discovery rate (log₂FC × –log₁₀[FDR]) from differential expression analysis. Gene Ontology (GO) gene sets were constructed by directly retrieving gene annotations from the *GO.db* package [48], specifically focusing on GO:0002376 (immune system process) and GO:0005102 (signaling receptor binding). Enrichment significance was assessed based on adjusted P-values and false discovery rate (FDR), allowing the identification of biological processes specifically enriched in the DPT cell population.

### TCR Clonotypes and Clonal Expansion Analysis

TCR sequencing data were analyzed using *VDJtools* to evaluate clonal expansion and repertoire diversity. Identical TCR clonotypes were defined as cells possessing the same paired TCR alpha and beta chain sequences. Sequence alignment and clonotype assignment enabled the quantification of clonotype frequency within both the DPT and SPT cell subsets. Clonal expansion was quantified by counting the number of cells assigned to each clonotype.

To examine temporal dynamics of DPT clonal expansion, we modeled changes in clonotype frequencies over time using the Generalized Additive Model (GAM). These dynamics were further stratified by disease severity to identify subset-specific expansion and contraction patterns. Data visualization was performed using custom R scripts and the *ggplot2* package (v3.5.1; Wickham, 2016).

Statistical comparisons of clonotype frequency and clonal expansion between the DPT and SPT subsets were conducted. The proportion of cells sharing identical clonotypes was calculated for each subset, and group differences were assessed using chi-squared and Fisher’s exact tests. A p-value of < 0.05 was considered statistically significant.

### Development of a Machine Learning Model for T cell Classification

A Random Forest classifier was developed to distinguish CD4⁺, CD8⁺, and DPT cells based on multi-gene transcriptional features rather than single-marker expression. The model development consisted of the following steps:

#### 1) Data Preparation

To address class imbalance during training, we applied an undersampling strategy. Specifically, two-thirds of the DPT cells and half of the predominant CD4⁺ and CD8⁺ cell populations were randomly selected for model training, while the remaining cells were reserved for testing. This approach ensured a more balanced distribution of cell types, facilitating effective model training and evaluation.

#### 2) Model Training and Hyperparameter Optimization

The classifier was based on a random forest algorithm, optimized through a comprehensive grid search involving 5,000 unique hyperparameter combinations. Key model parameters, including the number of decision trees (*ntree,* 100-2000), minimum node size (*nodesize,* 10-50), and the number of features per split (*mtry*, 5-30), were systematically tuned using 3-fold cross-validation (Fig. 2b).

#### 3) Handling Prediction Uncertainty

The Random Forest model outputs class probabilities for each prediction. To account for low-confidence classifications, cells for which the difference in probabilities between the top two predicted classes fell within the lowest 3% were labeled as “NA”. This conservative approach minimized erroneous classification in ambiguous cases.

#### 4) Model Evaluation

The model’s performance was assessed on the test set using standard evaluation metrics, including classification accuracy, precision, recall, and F1-score. These metrics provided a comprehensive overview of the model’s ability to distinguish between T cell subsets, with particular focus on the accurate identification of the relatively rare DPT cells. Confusion matrices were also generated to visualize classification performance across all classes.

### Feature importance analysis

To assess the relative contribution of individual genes to the classification task, feature importance scores were computed from the trained random forest model using the Gini importance metric, which quantifies the average gain in node purity provided by each feature across all decision trees in the ensemble. The importance scores were computed using the *getFeatureImportance* function from the *mlr* package (v2.19.1) [49] in R. This function calculates permutation-based importance by measuring the decrease in model performance when the values of each feature are randomly permuted, thereby capturing the impact of each gene on the overall predictive accuracy.

## Acknowledgments

This study was supported by grants from Korea University, Seoul, Republic of Korea (Grant Nos. K1913171 and K1507881). We thank Korea University Anam Hospital for providing computational infrastructure and administrative support. We are also grateful to our colleagues in the Department of Laboratory Medicine, Korea University Anam Hospital for their insightful discussions and constructive feedback.

We sincerely thank the healthy donors who generously provided blood samples for this study. We also acknowledge the dedicated efforts of our laboratory team in performing the single-cell RNA and TCR sequencing experiments. This work benefited from public datasets, including GSE158055, and we appreciate the researchers who made these valuable resources available to the scientific community.

## Funding

This study was supported by grants from Korea University, Seoul, Republic of Korea (Grant Nos. K1913171 and K1507881).

## Author contributions

**E.S.** designed and performed all bioinformatics analyses, including data processing, integrative analysis of scRNA-seq and scTCR-seq data, downstream analyses, and development of the supervised classification model for T cell subtype prediction. E.S. also drafted the manuscript. **S.G.Y.** contributed to single-cell sequencing experiments, prepared and managed IRB documentation, and reviewed the manuscript. **Y.J.C.** supervised the overall project, provided conceptual guidance, and reviewed the manuscript. All authors read and approved the final version of the manuscript.

## Data and code availability

The raw sequencing data generated in this study have been publicly deposited in the NCBI Sequence Read Archive (SRA) under accession number PRJNA1299552. Processed single-cell data and metadata are available through the NCBI Gene Expression Omnibus (GEO) under accession number GSE304309. The datasets generated and analyzed during this study are available from the corresponding author upon reasonable request. Publicly available COVID-19 single-cell data can be accessed through the NCBI Gene Expression Omnibus (GEO) under accession number GSE158055. The source code used to implement the random forest classifier is available at: https://github.com/shineunjhi/DPT-classifier.

## Competing interests

The authors declare no competing interests.

## Supporting information

**S1 Fig. Molecular features of DPT Cells in COVID-19 Patients.**

**S1 Table. Top 100 differentially expressed genes (DEGs) in each cluster of DPT cells compared to SPT cells.**

**S2 Table. Immune-related DEGs in each cluster identified in DPT cells compared to SPT cells.**

**S3 Table. Pathway enrichment of DEGs in CD4, CD8, and DPT cells.**

**S4 Table. Top-ranked features from the random forest classifier distinguishing T cell subtypes.**

## Notes

### Competing Interest Statement

The authors have declared that no competing interests exist.

## References

1. Rückert T, Lareau CA, Mashreghi M-F, Ludwig LS, Romagnani C. Clonal expansion and epigenetic inheritance of long-lasting NK cell memory. Nat Immunol. 2022;23: 1551–1563. doi:10.1038/s41590-022-01327-7

2. Sun L, Su Y, Jiao A, Wang X, Zhang B. T cells in health and disease. Sig Transduct Target Ther. 2023;8: 1–50. doi:10.1038/s41392-023-01471-y

3. Venturi V, Kedzierska K, Price DA, Doherty PC, Douek DC, Turner SJ, et al. Sharing of T cell receptors in antigen-specific responses is driven by convergent recombination. Proc Natl Acad Sci U S A. 2006;103: 18691–18696. doi:10.1073/pnas.0608907103

4. Blue ML, Daley JF, Levine H, Schlossman SF. Coexpression of T4 and T8 on peripheral blood T cells demonstrated by two-color fluorescence flow cytometry. J Immunol. 1985;134: 2281–2286.

5. Parel Y, Chizzolini C. CD4+ CD8+ double positive (DP) T cells in health and disease. Autoimmun Rev. 2004;3: 215–220. doi:10.1016/j.autrev.2003.09.001

6. Kitchen SG, Whitmire JK, Jones NR, Galic Z, Kitchen CMR, Ahmed R, et al. The CD4 molecule on CD8+ T lymphocytes directly enhances the immune response to viral and cellular antigens. Proc Natl Acad Sci U S A. 2005;102: 3794–3799. doi:10.1073/pnas.0406603102

7. Overgaard NH, Jung J-W, Steptoe RJ, Wells JW. CD4+/CD8+ double-positive T cells: more than just a developmental stage? J Leukoc Biol. 2015;97: 31–38. doi:10.1189/jlb.1RU0814-382

8. Schad SE, Chow A, Mangarin L, Pan H, Zhang J, Ceglia N, et al. Tumor-induced double positive T cells display distinct lineage commitment mechanisms and functions. J Exp Med. 2022;219: e20212169. doi:10.1084/jem.20212169

9. Yu ED, Wang H, da Silva Antunes R, Tian Y, Tippalagama R, Alahakoon SU, et al. A Population of CD4+CD8+ Double-Positive T Cells Associated with Risk of Plasma Leakage in Dengue Viral Infection. Viruses. 2022;14: 90. doi:10.3390/v14010090

10. Zhang H, Wang Y, Ma Y, Tang K, Zhang C, Wang M, et al. Increased CD4+CD8+ Double Positive T Cells during Hantaan Virus Infection. Viruses. 2022;14: 2243. doi:10.3390/v14102243

11. Menard LC, Fischer P, Kakrecha B, Linsley PS, Wambre E, Liu MC, et al. Renal Cell Carcinoma (RCC) Tumors Display Large Expansion of Double Positive (DP) CD4+CD8+ T Cells With Expression of Exhaustion Markers. Front Immunol. 2018;9: 2728. doi:10.3389/fimmu.2018.02728

12. Wu Y, Cai B, Feng W, Yang B, Huang Z, Zuo C, et al. Double positive CD4+CD8+ T cells: key suppressive role in the production of autoantibodies in systemic lupus erythematosus. Indian J Med Res. 2014;140: 513–519.

13. Marrero YT, Suárez VM, Abraham CMM, Hernández IC, Ramos EH, Domínguez GD, et al. Immunophenotypic characterization of double positive T lymphocytes in Cuban older adults. Exp Gerontol. 2021;152: 111450. doi:10.1016/j.exger.2021.111450

14. Hess NJ, Turicek DP, Riendeau J, McIlwain SJ, Contreras Guzman E, Nadiminti K, et al. Inflammatory CD4/CD8 double-positive human T cells arise from reactive CD8 T cells and are sufficient to mediate GVHD pathology. Science Advances. 2023;9: eadf0567. doi:10.1126/sciadv.adf0567

15. Mucida D, Husain MM, Muroi S, van Wijk F, Shinnakasu R, Naoe Y, et al. Transcriptional Reprogramming of Mature CD4+ T helper Cells generates distinct MHC class II-restricted Cytotoxic T Lymphocytes. Nat Immunol. 2013;14: 281–289. doi:10.1038/ni.2523

16. Alam MR, Akinyemi AO, Wang J, Howlader M, Farahani ME, Nur M, et al. CD4+CD8+ double-positive T cells in immune disorders and cancer: Prospects and hurdles in immunotherapy. Autoimmunity Reviews. 2025;24: 103757. doi:10.1016/j.autrev.2025.103757

17. Caccamo N, Joosten SA, Ottenhoff THM, Dieli F. Atypical Human Effector/Memory CD4+ T Cells With a Naive-Like Phenotype. Front Immunol. 2018;9. doi:10.3389/fimmu.2018.02832

18. Golubovskaya V, Wu L. Different Subsets of T Cells, Memory, Effector Functions, and CAR-T Immunotherapy. Cancers. 2016;8: 36. doi:10.3390/cancers8030036

19. Wang L, Wildt KF, Castro E, Xiong Y, Feigenbaum L, Tessarollo L, et al. The zinc finger transcription factor Zbtb7b represses CD8-lineage gene expression in peripheral CD4+ T cells. Immunity. 2008;29: 876–887. doi:10.1016/j.immuni.2008.09.019

20. Egawa T, Littman DR. ThPOK acts late in specification of the helper T cell lineage and suppresses Runx-mediated commitment to the cytotoxic T cell lineage. Nat Immunol. 2008;9: 1131–1139. doi:10.1038/ni.1652

21. Setoguchi R, Tachibana M, Naoe Y, Muroi S, Akiyama K, Tezuka C, et al. Repression of the Transcription Factor Th-POK by Runx Complexes in Cytotoxic T Cell Development. Science. 2008;319: 822–825. doi:10.1126/science.1151844

22. Feng D, Chen Y, Dai R, Bian S, Xue W, Zhu Y, et al. Chromatin organizer SATB1 controls the cell identity of CD4+ CD8+ double-positive thymocytes by regulating the activity of super-enhancers. Nature Communications. 2022;13: 5554. doi:10.1038/s41467-022-33333-6

23. Li C, Wu H, Guo L, Liu D, Yang S, Li S, et al. Single-cell transcriptomics reveals cellular heterogeneity and molecular stratification of cervical cancer. Commun Biol. 2022;5: 1–10. doi:10.1038/s42003-022-04142-w

24. Papalexi E, Satija R. Single-cell RNA sequencing to explore immune cell heterogeneity. Nat Rev Immunol. 2018;18: 35–45. doi:10.1038/nri.2017.76

25. Arora JK, James LK, Charoensawan V. Understanding and mitigating the impact of ambient mRNA contamination in single-cell RNA-sequencing analysis. [cited 22 Dec 2025]. doi:10.1371/journal.pone.0332440

26. Su M, Pan T, Chen Q-Z, Zhou W-W, Gong Y, Xu G, et al. Data analysis guidelines for single-cell RNA-seq in biomedical studies and clinical applications. Military Medical Research. 2022;9: 68. doi:10.1186/s40779-022-00434-8

27. Yamawaki TM, Lu DR, Ellwanger DC, Bhatt D, Manzanillo P, Arias V, et al. Systematic comparison of high-throughput single-cell RNA-seq methods for immune cell profiling. BMC Genomics. 2021;22: 66. doi:10.1186/s12864-020-07358-4

28. Quan F, Liang X, Cheng M, Yang H, Liu K, He S, et al. Annotation of cell types (ACT): a convenient web server for cell type annotation. Genome Medicine. 2023;15: 91. doi:10.1186/s13073-023-01249-5

29. Pasquini G, Rojo Arias JE, Schäfer P, Busskamp V. Automated methods for cell type annotation on scRNA-seq data. Comput Struct Biotechnol J. 2021;19: 961–969. doi:10.1016/j.csbj.2021.01.015

30. Heumos L, Schaar AC, Lance C, Litinetskaya A, Drost F, Zappia L, et al. Best practices for single-cell analysis across modalities. Nat Rev Genet. 2023;24: 550–572. doi:10.1038/s41576-023-00586-w

31. Ji Y, Lotfollahi M, Wolf FA, Theis FJ. Machine learning for perturbational single-cell omics. Cell Syst. 2021;12: 522–537. doi:10.1016/j.cels.2021.05.016

32. Moss P. The T cell immune response against SARS-CoV-2. Nat Immunol. 2022;23: 186–193. doi:10.1038/s41590-021-01122-w

33. Ren X, Wen W, Fan X, Hou W, Su B, Cai P, et al. COVID-19 immune features revealed by a large-scale single-cell transcriptome atlas. Cell. 2021;184: 1895–1913.e19. doi:10.1016/j.cell.2021.01.053

34. Moon J-S, Younis S, Ramadoss NS, Iyer R, Sheth K, Sharpe O, et al. Cytotoxic CD8+ T cells target citrullinated antigens in rheumatoid arthritis. Nat Commun. 2023;14: 319. doi:10.1038/s41467-022-35264-8

35. Drvar V, Ćurko-Cofek B, Karleuša L, Aralica M, Rogoznica M, Kehler T, et al. Granulysin expression and granulysin-mediated apoptosis in the peripheral blood of osteoarthritis patients. Biomed Rep. 2022;16: 44. doi:10.3892/br.2022.1527

36. Wohlfert EA, Grainger JR, Bouladoux N, Konkel JE, Oldenhove G, Ribeiro CH, et al. GATA3 controls Foxp3+ regulatory T cell fate during inflammation in mice. J Clin Invest. 2011;121: 4503–4515. doi:10.1172/JCI57456

37. Rudra D, deRoos P, Chaudhry A, Niec R, Arvey A, Samstein RM, et al. Transcription factor Foxp3 and its protein partners form a complex regulatory network. Nat Immunol. 2012;13: 1010–1019. doi:10.1038/ni.2402

38. Mantel P-Y, Kuipers H, Boyman O, Rhyner C, Ouaked N, Rückert B, et al. GATA3-Driven Th2 Responses Inhibit TGF-β1–Induced FOXP3 Expression and the Formation of Regulatory T Cells. [cited 13 Jun 2025]. doi:10.1371/journal.pbio.0050329

39. Baudu T, Parratte C, Perez V, Ancion M, Millevoi S, Hervouet E, et al. The NMD Pathway Regulates GABARAPL1 mRNA during the EMT. Biomedicines. 2021;9: 1302. doi:10.3390/biomedicines9101302

40. Schoppmeyer R, Zhao R, Cheng H, Hamed M, Liu C, Zhou X, et al. Human profilin 1 is a negative regulator of CTL mediated cell-killing and migration. European Journal of Immunology. 2017;47: 1562–1572. doi:10.1002/eji.201747124

41. Hu Z-W, Chen L, Ma R-Q, Wei F-Q, Wen Y-H, Zeng X-L, et al. Comprehensive analysis of ferritin subunits expression and positive correlations with tumor-associated macrophages and T regulatory cells infiltration in most solid tumors. Aging (Albany NY). 2021;13: 11491–11506. doi:10.18632/aging.202841

42. Starr TK, Jameson SC, Hogquist KA. Positive and negative selection of T cells. Annu Rev Immunol. 2003;21: 139–176. doi:10.1146/annurev.immunol.21.120601.141107

43. Ng SS, De Labastida Rivera F, Yan J, Corvino D, Das I, Zhang P, et al. The NK cell granule protein NKG7 regulates cytotoxic granule exocytosis and inflammation. Nature Immunology. 2020;21: 1205–1218. doi:10.1038/s41590-020-0758-6

44. Xu H, Lin S, Zhou Z, Li D, Zhang X, Yu M, et al. New genetic and epigenetic insights into the chemokine system: the latest discoveries aiding progression toward precision medicine. Cellular & Molecular Immunology. 2023;20: 739–776. doi:10.1038/s41423-023-01032-x

45. McGovern KE, Wilson EH. Role of Chemokines and Trafficking of Immune Cells in Parasitic Infections. Curr Immunol Rev. 2013;9: 157–168. doi:10.2174/1573395509666131217000000

46. Hao Y, Hao S, Andersen-Nissen E, Mauck WM, Zheng S, Butler A, et al. Integrated analysis of multimodal single-cell data. Cell. 2021;184: 3573–3587.e29. doi:10.1016/j.cell.2021.04.048

47. Eide PW, Bruun J, Lothe RA, Sveen A. CMScaller: an R package for consensus molecular subtyping of colorectal cancer pre-clinical models. Scientific Reports. 2017;7: 16618. doi:10.1038/s41598-017-16747-x

48. Ashburner M, Ball CA, Blake JA, Botstein D, Butler H, Cherry JM, et al. Gene Ontology: tool for the unification of biology. Nature Genetics. 2000;25: 25–29. doi:10.1038/75556

49. Bischl B, Lang M, Kotthoff L, Schiffner J, Richter J, Studerus E, et al. mlr: Machine Learning in R. Journal of Machine Learning Research. 2016;17: 1–5.

